# Different populations of midbrain neurons control the production of distinct acoustic categories of vocalization

**DOI:** 10.1101/2023.09.15.557179

**Authors:** Patryk Ziobro, Yena Woo, Zichen He, Katherine Tschida

## Abstract

Animals produce different acoustic types of vocalizations in different behavioral contexts. The midbrain periaqueductal gray (PAG) is obligatory for vocal production (Jürgens, 1994, 2002, 2009), but how midbrain circuits are organized to control the production of different vocalization types remains unknown. To test whether a single population of midbrain neurons regulates the production of different vocalization types, we applied intersectional methods to selectively ablate a population of midbrain neurons required for the production of ultrasonic vocalizations (USVs) in mice. We find that although ablation of these PAG-USV neurons blocks USV production in both males and females, these neurons are not required for the production of squeaks. Our findings provide strong evidence that distinct populations of midbrain neurons control the production of different acoustic categories of vocalizations.

## Results

A fundamental feature of vocal communication is that animals produce vocalizations with different acoustic features in different behavioral contexts (i.e., contact calls, territorial calls, courtship calls, etc.). The midbrain periaqueductal gray (PAG) plays an obligatory role in vocalization (Adametz and O’Leary, 1959; Skultety, 1962, 1965; Jürgens and Pratt, 1979; Jürgens, 1994, 2002, 2009; Esposito et al., 1999; Kittelberger et al., 2006), but how PAG circuits are organized to control the production of different vocalization types remains unknown. On the one hand, studies have found that partial lesions of the PAG abolish the production of some vocalization types while leaving others intact (Kelly et al., 1946; Skultety, 1962; Newman and MacLean, 1982; Jürgens, 1994), suggesting that different populations of PAG neurons might control the production of different vocalization types. On the other hand, electrophysiological recordings have revealed individual PAG neurons that increase their activity during the production of multiple vocalization types (Larson and Kistler, 1984, 1986; Düsterhöft et al., 2000), suggesting that some PAG neurons may contribute to the production of multiple types of vocalizations.

In the past, the inability to selectively manipulate vocalization-related PAG neurons made it challenging to resolve these conflicting findings. The recent identification of specialized PAG neurons whose activity is required for the production of male mouse courtship ultrasonic vocalizations (PAG-USV neurons; Tschida et al., 2019) opens the door to understanding whether a single population of midbrain neurons controls the production of multiple vocalization types. Mice produce USVs during same-sex and opposite-sex social interactions (Whitney et al., 1974; Nyby et al., 1979; Nyby, 1983; Moles et al., 2007; Neunuebel et al., 2015; Warren et al., 2018, 2020; Sangiamo et al., 2020) and also produce human-audible harmonic vocalizations (squeaks) in aversive contexts (Gourbal et al., 2004; Grimsley et al., 2013, 2016). In the current study, we tested whether selective ablation of PAG-USV neurons affects the production of squeaks.

### Ablation of PAG-USV neurons blocks USV production in both male and female mice

To ablate PAG-USV neurons in male and female mice, we used the TRAP2 activity-dependent labeling strategy (Allen et al., 2017; DeNardo et al., 2019). Because this approach differs from the activity-dependent labeling method used previously to manipulate PAG-USV neurons (Tschida et al., 2019), we first confirmed that TRAP2-mediated ablation of PAG-USV neurons blocks USV production. In addition, because previous work tested the effects of PAG-USV manipulations on USV production in males only, here we also tested the effects of PAG-USV ablation on female USV production.

Briefly, the PAG of male and female TRAP2;Ai14 mice was injected bilaterally with a virus driving the Cre-dependent expression of caspase (AAV-FLEX-taCasp3-TEVp; Fig. 1A). Mice were subsequently returned to group-housing with their same-sex siblings for 11 days and were single-housed for 3 days prior to the first behavioral measurements to promote high levels of social interaction and USV production (Zhao et al., 2021). Two weeks following viral injections (day 14), male and female subject mice were given 30-minute social encounters with a novel, group-housed female in their home cage, a context which elicits high rates of USV production in both males and females (Zhao et al., 2021). Following these social encounters, mice received I.P. injections of 4-hydroxytamoxifen (4-OHT), which enables the transient expression of Cre recombinase in recently active neurons, hence permitting the expression of caspase in PAG-USV neurons. Ten days later, male and female subjects were given a second 30-minute social encounter with a novel, group-housed female, and rates of USV production and non-vocal social behaviors were compared between the pre-4-OHT and post-4-OHT behavior sessions.

**Figure 1.**
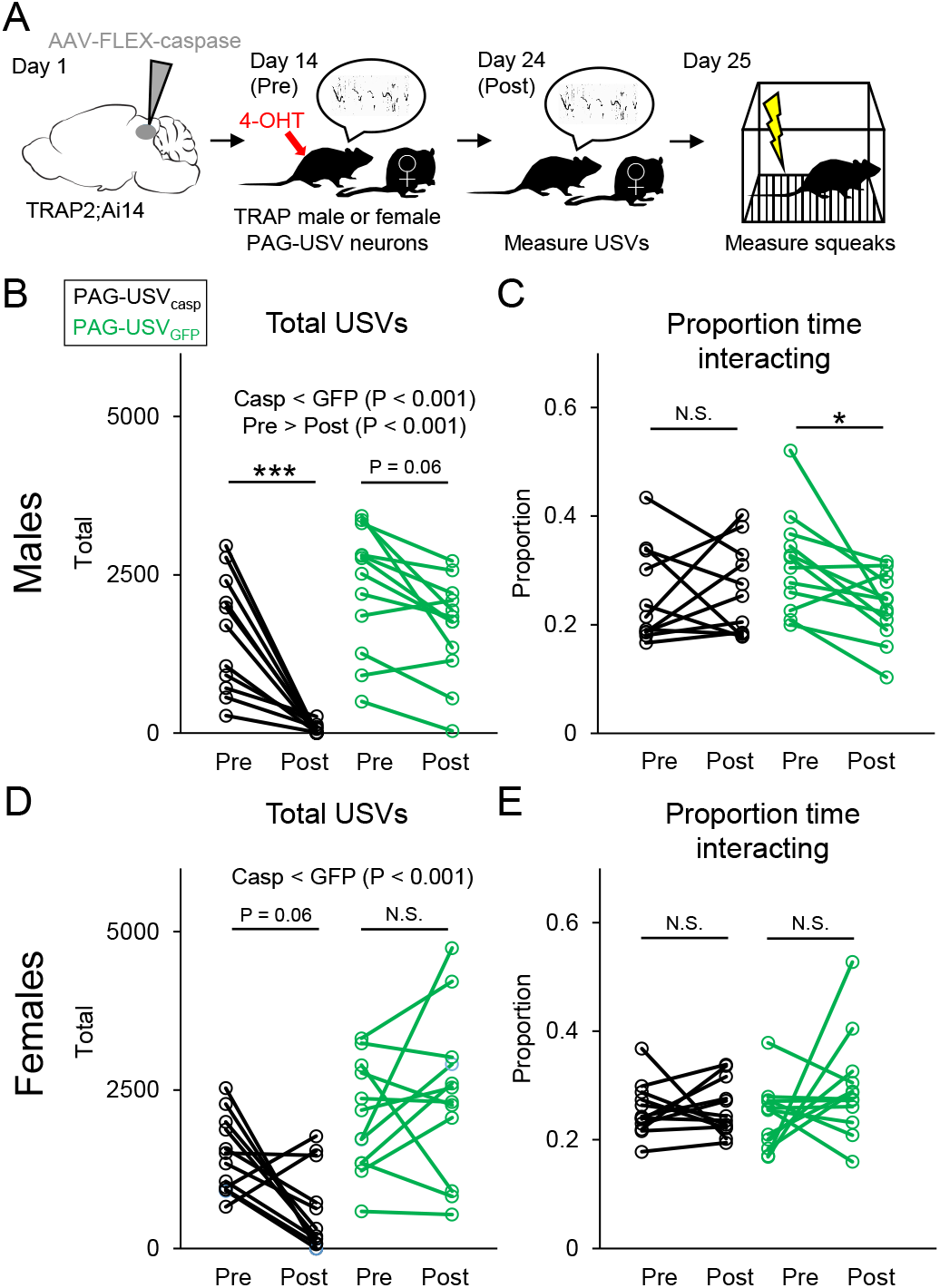
PAG-USV neurons are required for USV production in male and female mice. (A) Schematic shows experimental timeline for TRAP2-mediated ablation of PAG-USV neurons and behavioral measurements. (B) Total USVs produced by experimental (PAG-USV_casp_, black points, N = 11) and control (PAG-USV_GFP_, green points, N = 12) male mice during social interactions with females are shown before and after 4-OHT treatment. (C) Same as (B), for proportion time subject males spent interacting with females before and after 4-OHT treatment. (D) Total USVs produced by experimental (PAG-USV_casp_, black points, N = 12) and control (PAG-USV_GFP_, green points, N = 12) female mice during social interactions with females are shown before and after 4-OHT treatment. (E) Same as (D), for proportion time subject females spent interacting with females before and after 4-OHT treatment. In (B-E), statistically significant main effects are indicated in text centered above plots, and results of post-hoc pairwise comparisons are indicated above the relevant comparisons. See Table S1 for complete statistical details.

Using this approach, we first measured the effects of PAG-USV ablation on male USV production. Previous studies using microphone arrays to localize and assign USVs to individual mice reported that males produce ∼85% of total USVs recorded during male-female interactions (Neunuebel et al., 2015; Warren et al., 2021). Based on these findings, we assumed that the majority of USVs recorded during our male-female interactions were produced by the male. Consistent with previous work (Tschida et al., 2019), ablation of PAG-USV neurons dramatically reduced USV production in male mice (Fig. 1B, black symbols; N = 11 PAG-USV_casp_ males; pre-4-OHT mean USVs = 1582 ± 928; post-4-OHT mean USVs = 53 ± 84; p < 0.001, two-way ANOVA with repeated measures on one factor and post-hoc Tukey’s HSD tests; see Table S1 for complete statistical details). In contrast to USV production, the amount of time males spent engaged in non-vocal interactions with females was not affected by PAG-USV ablation (Fig. 1C, black symbols; mean proportion time interacting with female pre-4-OHT = 0.25 ± 0.09; post-4-OHT = 0.26 ± 0.08; p = 0.77, two-way ANOVA with repeated measures on one factor and post-hoc Tukey’s HSD tests). In control TRAP2;Ai14 males with GFP expressed in PAG-USV neurons, USV rates tended to decline (p = 0.06) and time spent interacting with females declined significantly (p = 0.04; Fig. 1B-C, green symbols; N = 12 PAG-USV_GFP_ males; pre-4-OHT mean USVs = 2309 ± 985 USVs; post-4-OHT mean USVs = 1664 ± 787 USVs; pre-4-OHT mean proportion time interacting with female = 0.31 ± 0.09; post-4-OHT mean proportion time interacting with female = 0.24 ± 0.06). To more clearly quantify the magnitude of change in USV rates over time for each group, we calculated the change in USV rates for each male (change in USV rate= total post-4-OHT USVs / total pre-4-OHT USVs). A comparison of these values showed that males that underwent PAG-USV ablation exhibited a much stronger decrease in USV production than control males (mean post-4-OHT / pre-4-OHT USVs for PAG-USV_casp_ males = 0.07 ± 0.12; mean post-4-OHT / pre-4-OHT USVs for PAG-USV_GFP_ males = 0.71 ± 0.32; p < 0.001, t-test). In summary, we conclude that TRAP2-mediated ablation of PAG-USV neurons reduces USV production in male mice without affecting time spent engaged in social interactions with females, consistent with prior work (Tschida et al., 2019).

We next measured the effects of PAG-USV ablation on female USV production. Previous studies found that females vocalize at the highest rates during same-sex interactions (Moles et al., 2007; Warren et al., 2020; Zhao et al., 2021) and that both females in a pair produce USVs during same-sex interactions (Warren et al., 2020). Given these findings, we reasoned that we should see reduced (but non-zero) USV rates during female-female interactions in which one partner has undergone ablation of PAG-USV neurons. In line with this idea, we observed that ablation of PAG-USV neurons tended to reduce USV production during female-female interactions (Fig. 1D, black symbols; N = 12 PAG-USV_casp_ females; pre-4-OHT mean USVs =1468 ± 599 USVs; post-4-OHT mean USVs = 579 ± 657; p = 0.06, two-way ANOVA with repeated measures on one factor and post-hoc Tukey’s HSD tests). Rates of USVs did not change over time in pairs containing a PAG-USV_GFP_ female (Fig. 1D, green symbols; N = 12 GFP females; pre-4-OHT mean USVs = 2060 ± 873 USVs; post-4-OHT mean USVs = 2410 ± 1273 USVs; p = 0.77). We next reasoned that we should see near-zero USV rates in female-female interactions in which both partners have undergone ablation of PAG-USV neurons. Consistent with this idea, we found that USV rates were near-zero during recordings from pairs of females that had both undergone ablation of PAG-USV neurons (Fig. S1, right column; mean USVs = 49 ± 36; N = 8 trials from pairs made up of N = 7 total PAG-USV_casp_ females). These USV rates were significantly lower than USV rates recorded from control pairs containing 1 PAG-USV_GFP_ female and 1 unmanipulated female (Fig.S1, left column; p < 0.001; one-way ANOVA with post-hoc Tukey HSD tests). Ablation of PAG-USV neurons had no effect on the amount of time females spent engaged in non-vocal interactions with female partners (Fig. 1E, black symbols; pre-4-OHT mean proportion time interacting with female partner = 0.25 ± 0.05; post-4-OHT = 0.26 ± 0.05), and control PAG-USV_GFP_ females also showed no change in social interaction time following 4-OHT treatment (Fig. 1E, green symbols; pre-4-OHT mean proportion time interacting with female partner = 0.24 ± 0.06; post-4-OHT = 0.30 ± 0.10; two-way ANOVA with repeated measures on one factor; p > 0.05 for main effects and interaction). In summary, we conclude that ablation of PAG-USV neurons blocks USV production in both female and male mice without affecting time spent engaged in social interactions with female partners.

### PAG-USV neurons are not required for the production of squeaks

With these strategies in hand, we next asked whether ablation of PAG-USV neurons in males and females blocks the production of squeaks. The same PAG-USV_casp_ and PAG-USV_GFP_ males and females that were tested for USV production (Figs. 1, S1) were subjected to a mild footshock paradigm the day following their second social interaction (day 25, Fig. 1A; each mouse received 10 footshocks delivered over 5 minutes, see Methods). We selected a footshock intensity (0.5 mA) that reliably elicited squeaks in control PAG-USV_GFP_ males and females (Fig. 2A, right; 116/120 and 119/120 footshocks elicited squeaks in N = 12 control males and N = 12 control females, respectively). In contrast to the pronounced effects of PAG-USV ablation on USV production, the ablation of PAG-USV neurons did not block the production of squeaks, and both male and female PAG-USV_casp_ mice produced squeaks reliably in response to footshock (Fig. 2A, left; 120/120 and 120/120 footshocks elicited squeaks in N =12 PAG-USV_casp_ males and N = 12 PAG-USV_casp_ females, respectively; two-way ANOVA, p > 0.05 for main effects of sex, group, and interaction). We conclude that the activity of PAG-USV neurons is not required for the production of squeaks.

**Figure 2.**
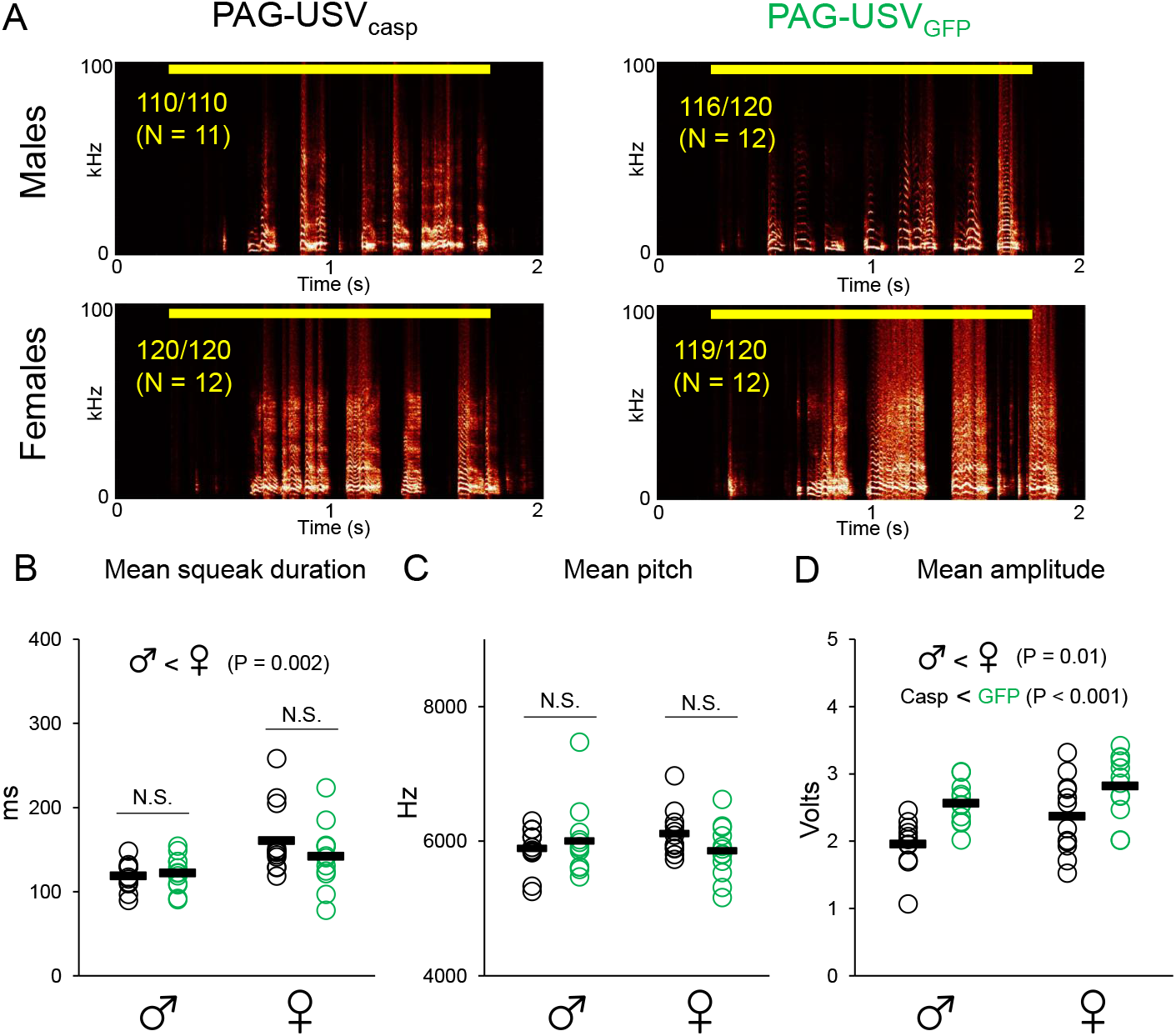
PAG-USV neurons are not required for the production of squeaks. (A) Representative spectrograms show squeaks produced in response to footshocks for experimental (PAG-USV_casp_) and control (PAG-USV_GFP_) male and female mice. Yellow bar at the top of each spectrogram indicates timing of footshock. Yellow text indicates proportion of total trials across all mice in each group in which footshocks elicited squeaks. (B) Mean squeak duration is plotted for male (left) and female (right) mice from experimental (PAG-USV_casp_) and control (PAG-USV_GFP_) groups. (C) Same as (B), for mean squeak pitch. (D) Same as (B), for mean squeak amplitude. In (B-D), statistically significant main effects are indicated in text centered above plots, and results of post-hoc pairwise comparisons are indicated above the relevant comparisons. See Table S1 for complete statistical details.

Although the activity of PAG-USV neurons is not required for the production of squeaks, it remains possible that the activity of these neurons plays a more subtle role in modulating the acoustic features of squeaks. We examined this possibility by quantifying the acoustic features of squeaks produced by PAG-USV_casp_ and PAG-USV_GFP_ mice (see Methods). Although mean squeak duration was greater in females than in males, there was no effect of PAG-USV ablation on squeak duration (Fig. 2B; two-way ANOVA, p = 0.002 for main effect of sex, p > 0.05 for main effect of group and interaction). Ablation of PAG-USV neurons also had no effect on the mean pitch of squeaks, and mean pitch did not differ significantly between females and males (Fig. 2C; two-way ANOVA, p > 0.05 for main effects and interaction). Interestingly, although female mice on average produced louder squeaks than males (Fig. 2D; two-way ANOVA, p = 0.02 for main effect of sex), overall mice that underwent ablation of PAG-USV neurons produced squeaks that were quieter than those produced by control PAG-USV_GFP_ mice (Fig. 2D, p < 0.001 for main effect of group; p = 0.54 for interaction). One possibility is that although PAG-USV neurons are not required for squeak production, the activity of these neurons regulates the amplitude of non-USV vocalizations. Another possibility is that the effects on squeak amplitude can be attributed to the ablation of non-PAG-USV neurons that increase their activity around the time of 4-OHT treatment. Although behavior sessions are designed to maximize USV production, mice also engage in various non-vocal behaviors and experiences that may increase activity in non-PAG-USV neuronal populations. For example, PAG neurons related to fear responses and nociception may increase their activity as mice are handled by the investigator and receive an IP injection of 4-OHT. To test whether we could recapitulate the effects on squeak amplitude following ablation of PAG neurons that are recruited by any such non-vocal/non-social experiences, we performed control experiments in which caudolateral PAG neurons were ablated in mice that were placed alone in their homecage inside the recording chamber for 30 minutes and then given an IP injection of 4-OHT (Fig. 3A, N = 5 PAG-control_casp_ males). Ablation of PAG-control_casp_ neurons did not reduce rates of USV production (Fig. 3B; p = 0.16, paired t-test), nor did it affect the acoustic features of USVs (pre-4-OHT mean duration = 47.0 ± 9.1 ms; post-4-OHT mean duration = 49.7 ± 9.0 ms; pre-4-OHT mean pitch = 69.2 ± 4.4 kHz; post-4-OHT mean pitch = 69.8 ± 3.3 kHz; pre-4-OHT mean dB relative to background = 14.9 ± 3.9 dB; post-4-OHT mean dB relative to background = 11.4 ± 1.0 dB; paired t-tests, p > 0.05 for all). Similarly, ablation of PAG-control_casp_ neurons did affect the production of squeaks (Fig. 3C; 48/50 footshocks elicited squeaks in N = 5 PAG-control_casp_ males; one-way ANOVA), mean squeak duration (Fig. 3D), or mean squeak pitch (Fig. 3E; one-way ANOVAs, p > 0.05 for all). Notably, PAG-control_casp_ males produced squeaks that were significantly lower in amplitude than squeaks produced by both PAG-USV_casp_ and PAG-USV_GFP_ males (Fig. 3F; one-way ANOVA, p < 0.001 for all pair-wise post-hoc comparisons). This finding is consistent with the idea that PAG-USV neurons regulate neither the production nor the acoustic features of squeaks, and that reduced squeak amplitude in PAG-USV_casp_ mice can be attributed to the ablation of non-PAG-USV neurons that increase their activity during or shortly following the behavior session and 4-OHT treatment.

**Figure 3.**
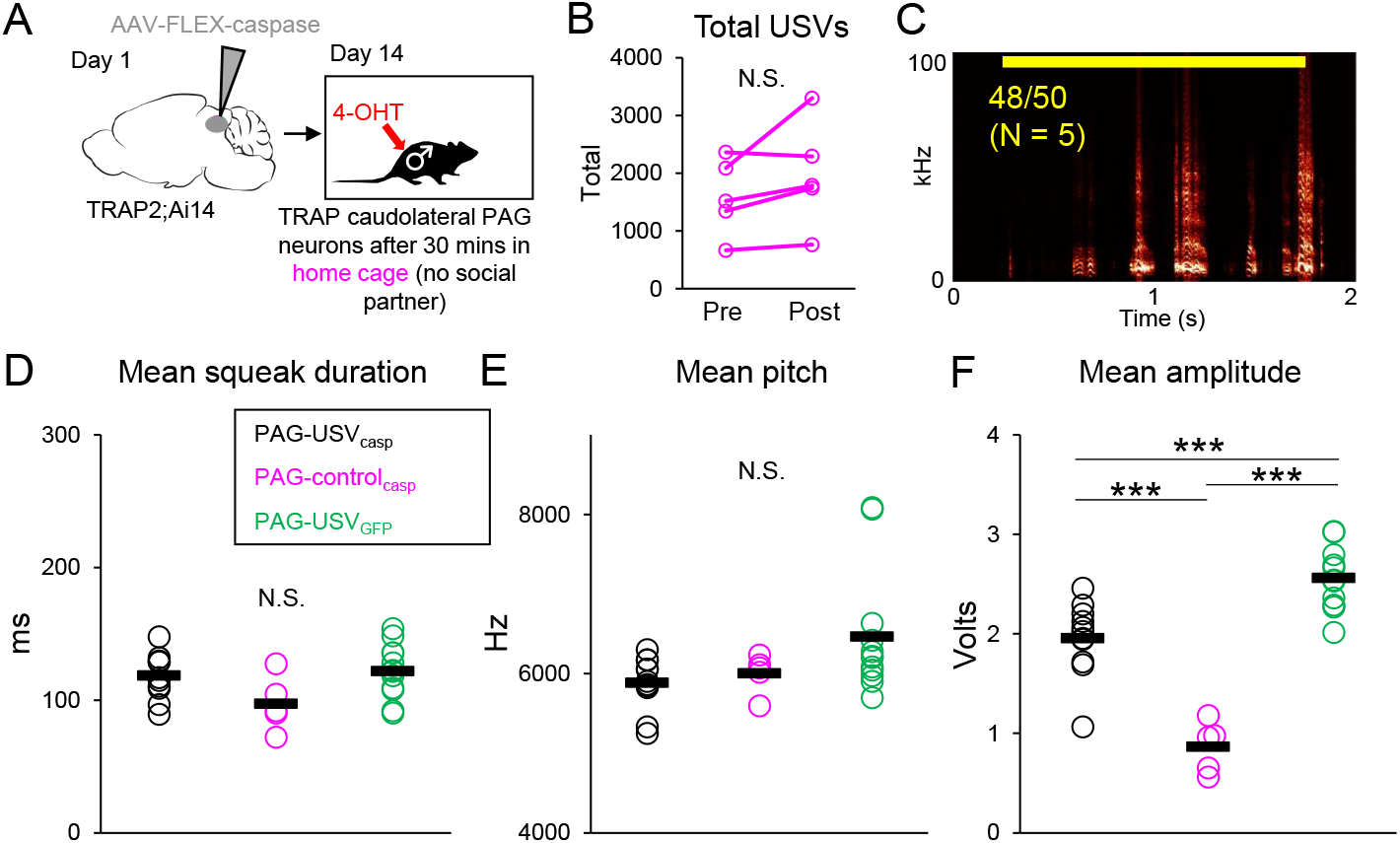
Effects of ablation of control (non-PAG-USV) caudolateral PAG neurons on production of USVs and squeaks in male mice. (A) Schematic shows experimental timeline for TRAP2-mediated ablation of control, non-PAG-USV neurons in male mice. (B) Total USVs produced by PAG-control_casp_ males (N = 5) during interactions with females are shown before and after 4-OHT treatment. (C) Representative spectrogram shows squeaks produced by a PAG-control_casp_ male in response to footshock. Yellow bar at the top of the spectrogram indicates timing of footshock. Yellow text indicates proportion of total trials across N = 5 PAG-control_casp_ males in which footshock elicited squeaks. (D) Mean squeak duration is plotted for PAG-USV_casp_ males (black points, N = 11), PAG-control_casp_ males (magenta points, N = 5), and PAG-USV_GFP_ males (green points, N = 12). (E) Same as (D), for mean squeak pitch. (F) Same as (D), for mean squeak amplitude.

Finally, we tested whether ablation of PAG-USV neurons in female mice affects the production of squeaks produced in a social context, during courtship interactions with males. We found that 9 out of 9 PAG-USV_casp_ females produced squeaks during interactions with males (Fig. S2), indicating that ablation of PAG-USV neurons does not block the production of squeaks in social contexts. In summary, these results provide strong support for the idea that PAG-USV neurons do not regulate the production of squeaks and that distinct populations of neurons control the production of different acoustic types of vocalizations.

## Discussion

In the current study, we confirm previous work showing that the activity of a specialized population of PAG-USV neurons is essential for USV production in male mice (Tschida et al., 2019), and we extend these findings to show that ablation of PAG-USV neurons also impairs USV production in females. In contrast to the robust effects on USV production, ablation of PAG-USV neurons did not block the production of squeaks and did not affect the duration or pitch of squeaks produced by males and females in response to footshock. Although we observed a significant decrease in squeak amplitude following ablation of PAG-USV neurons, these effects were not specific to PAG-USV neuronal ablation and were also observed following ablation of control (non-PAG-USV) neurons in the caudolateral PAG. The fact that ablation of such control neurons failed to affect rates of USV production or USV acoustic features underscores the selective role of PAG-USV neurons in regulating USV production. Taken together, our findings indicate that distinct populations of neurons control the production of USVs and squeaks.

Neurons important for the production of squeaks remain to be identified. Although we do not know for certain that such neurons reside within the PAG, a wealth of previous work in non-murine species has shown that extensive lesions of the PAG cause complete mutism (Jürgens, 1994, 2002, 2009). However, given the well characterized roles of the PAG in regulating fear behaviors and nociception and the fact that mice produce squeaks almost exclusively when in fear or when in pain, the use of activity-dependent labeling approaches to identify such neurons presents a particular challenge. A recent study in mice found that pharmacological inhibition of the dorsolateral PAG abolished the production of squeaks, but the concomitant effects on the expression of fear behaviors make it difficult to know whether this region of the PAG directly or indirectly regulates squeak production (Ruat et al., 2022). Given the well-known role of the hindbrain nucleus retroambiguus (RAm) and PAG projections to RAm in regulating vocal production (Holstege, 1989; Zhang et al., 1992; Shiba et al., 1997; Vanderhorst et al., 2000; VanderHorst et al., 2001; Jürgens, 2002, 2009; Wild et al., 2009; Tschida et al., 2019), an intersectional approach that restricts activity-dependent labeling to RAm-projecting PAG neurons may provide a useful future strategy to identify and characterize midbrain neurons that regulate the production of squeaks. It would be of great interest to characterize the axonal projections of PAG neurons important for squeak production and to compare them to those of PAG-USV neurons (Tschida et al., 2019), to begin to understand how differences in the activity of midbrain-to-hindbrain circuits underlie the production of distinct acoustic types of vocalization.

In addition to being produced in relatively non-overlapping behavioral contexts, USVs and squeaks are also produced via distinct sound production mechanisms (i.e., vibration of the folds to generate squeaks vs. an intralaryngeal whistle mechanism to generate USVs (Roberts, 1975; Riede, 2011, 2013; Mahrt et al., 2016; Riede et al., 2017 ; Håkansson et al., 2022). In light of these differences, one interpretation of our results is that while distinct populations of midbrain neurons regulate the production of sonic and ultrasonic vocalizations, a single population may regulate the production of acoustic types of vocalizations that share a sound production mechanism but differ in their contextual usage. For example, in rats, does a single population of PAG-USV neurons control the production of both 50 kHz and 22 kHz USVs? Another possibility is that distinct populations of midbrain neurons regulate the production of acoustic types of vocalizations that differ in their contextual usage, even if these acoustic types have similar sound production mechanisms. Our results provide strong support for the idea that different populations of midbrain neurons regulate the production of distinct acoustic types of vocalizations, and these findings form an important foundation to understand how midbrain circuits are organized to regulate vocal communication across diverse species with diverse vocal repertoires.

## Acknowledgements

Thanks to Xin Zhao, Da-Jiang Zheng, David Smith, and Thom Cleland for providing feedback on draft versions of this manuscript. Thanks also to Frank Drake and other CARE staff for their excellent mouse husbandry.

## Author contributions

P.Z. and K.A.T. designed the experiments. P.Z. conducted the experiments. P.Z., Y.W., Z.H., and K.A.T. analyzed the data. P.Z. and K.A.T. wrote the manuscript, and all authors approved the final version.

## Materials and Methods

### Resource availability

Further information and requests for resources should be directed to and will be fulfilled by the lead contact, Katherine Tschida. No materials were generated for this study.

### Ethics Statement

All experiments and procedures were conducted according to protocols approved by the Cornell University Institutional Animal Care and Use Committee (protocol #2020-001).

### Subject Details

Male and female TRAP2;Ai14 mice were housed with their siblings and both parents until weaning at postnatal day 21. TRAP;A14 mice were generated by crossing TRAP2 (Jackson Laboratories, 030323) with Ai14 (Jackson Laboratories, 007914). Mice were kept on a 12h:12h reversed light/dark cycle and given ad libitum food and water for the duration of the experiment.

### Viral injections

The following viruses were used: AAV2/5-ef1alpha-FLEX-taCasp3-TEVp (Addgene, 45580) and AAV2/1-CAG-FLEX-EGFP-WPRE (Addgene, 51502). The final injection coordinates for caudolateral PAG were: AP = -4.7 mm, ML = 0.5 mm, DV = 1.75 mm. Viruses were pressure-injected with a Nanoject III (Drummond) at a rate of 5 nL every 20 seconds. A total volume of 250 nl of virus was injected into each side of the PAG.

### Drug preparation

4-hydroxytamoxifen (4-OHT, HelloBio, HB6040) was dissolved at 20 mg/mL in ethanol by shaking at 37°C and was then aliquoted (75 uL) and stored at -20°C. Before use, 4-OHT was redissolved in ethanol by shaking at 37°C and filtered corn oil was added (Sigma, C8267, 150 uL). Ethanol was then evaporated by vacuum under centrifugation to give a final concentration of 10 mg/mL, and 4-OHT solution was used on the same day it was prepared.

### Study design

To express either caspase or GFP in PAG-USV neurons, adult (>8 weeks) male and female subject mice first received bilateral injections of virus (caspase virus for PAG-USV_casp_ mice, GFP virus for PAG-USV_GFP_ mice) into the caudolateral PAG (day 0). Eleven days later, subject mice were single-housed for 3 days. On day 14, subject mice were given 30-minute social interactions with a novel, group-housed female visitor to elicit USV production (see below for details of vocal and non-vocal behavior recording and analysis). Following the social interaction, subject mice received I.P. injections of 4-OHT (50 mg/kg) to enable expression of viral transgenes in PAG-USV neurons. On day 24, subject mice were given a second 30-minute social encounter with a novel, group-housed female to measure effects of viral expression on USV production and non-vocal social behavior. On day 25, mice were subjected to a mild footshock paradigm to measure effects of viral expression on the production and acoustic features of squeaks.

To express caspase in control (non-PAG-USV) neurons of the caudolateral PAG, adult male mice first received bilateral injections of caspase virus into the caudolateral PAG (day 0). Eleven days later, males were single-housed for 3 days. On day 13, males were given 30-minute social interactions with a novel, group-housed female visitor to elicit and measure baseline USV production. On day 14, males in their home cages were placed inside the recording chamber but not given a social partner. After 30 minutes, males received I.P. injections of 4-OHT (50 mg/kg) to enable expression of caspase in control PAG neurons. On day 24, these PAG-control_casp_ males were given a second 30-minute social encounter with a novel, group-housed female to measure effects of viral expression on USV production and non-vocal social behavior. On day 25, PAG-control_casp_ males were subjected to a mild footshock paradigm to measure effects of viral expression on the production and acoustic features of squeaks.

### USV recording and analyses

To elicit USVs, single-housed male and female subject mice were given a 30-minute social experience with a novel, group-housed female visitor in their home cage. Home cages were placed in a sound-attenuating chamber (Med Associates) and fitted with custom lids that allowed microphone access and permitted audio (and video) recordings. USVs were recorded with an ultrasonic microphone (Avisoft, CMPA/CM16), amplified (Presonus TubePreV2), and digitized at 250 kHz (Spike 7, CED). USVs were detected with custom Matlab codes (Tschida et al., 2019) using the following parameters: mean frequency > 45 KHz; spectral purity > 0.3; spectral discontinuity < 0.85; minimum USV duration = 5 ms; minimum inter-syllable interval = 30 ms). The duration of each USV was calculated as the difference between USV onset and offset. The mean pitch of each USV was determined by calculating the dominant frequency at each time bin of the USV and then averaging across the syllable (Tschida et al., 2019). The amplitude of each USV was defined as the bandpower from 30-125 kHz, converted to dB and measured relative to periods of quiet background noise recorded in the same trial (Tschida et al., 2019).

### Analyses of non-vocal social behaviors

Non-vocal behavior was recorded in each trial with a webcam (Logitech). BORIS software was used by trained observers to score and record non-vocal behaviors from video recordings of pairs of interacting mice. The following behaviors were recorded: resident-initiated social investigation (sniffing, following, or chasing), visitor-initiated social investigation, mutual social investigation, and resident-initiated mounting (no instances of visitor-initiated mounting were observed in our dataset).

### Footshock delivery

Subject mice were placed in a footshock chamber (Med Associates) inside of a sound attenuating chamber (Med Associates) equipped with an ultrasonic microphone (Avisoft) and a webcam (Logitech). A mild (0.5 mA) shock was delivered once every 30 seconds, for a total of 10 footshocks (trial duration = 5 minutes). Audio and video recordings were conducted as described above for USVs.

### Analysis of squeak acoustic features

A custom Matlab code was used that allowed trained users to manually annotate the onsets and offsets of individuals squeaks from spectrograms created from Spike2 audio recordings. The duration of individual squeaks was then calculated from these annotations. Another custom Matlab code was used to calculate the mean pitch (mean dominant frequency) and the mean amplitude of each squeak. Because the microphone was placed near the top of the footshock chamber, audio clipping occurred during portions of squeaks in many trials. We therefore report mean squeak amplitudes in volts, directly measured as the mean amplitude of the audio waveform recorded in Spike2 during manually annotated squeak times.

### Post-hoc visualization of viral labeling

Mice were deeply anesthetized with isoflurane and transcardially perfused with ice-cold 4% paraformaldehyde in 0.1 M phosphate buffer, pH 7.4 (4% PFA). Dissected brains were post-fixed overnight in 4% PFA at 4°C, cryoprotected in a 30% sucrose solution in PBS at 4°C for 48 hours, frozen in embedding medium (Surgipath, VWR), and stored at -80°C until sectioning. Brains were cut into 80 um coronal sections on a cryostat, rinsed 3 × 10 minutes in PBS, and processed at 4°C with NeuroTrace (1:500, Invitrogen) in PBS containing 0.3% Triton-X. Tissue sections were rinsed again 3 × 10 minutes in PBS, mounted on slides, and coverslipped with Fluoromount-G (Southern Biotech). After drying, slides were imaged with a 10x objective on a Zeiss LSM900 confocal laser scanning microscope. Because expression of the caspase virus cannot be directly visualized by looking for expression of a fluorescent tag, we took the following approach to assess the spread of the caspase virus. In TRAP2;Ai14 mice that are treated with 4-OHT following a social encounter, neurons throughout the brain that upregulate Fos during the social encounter will be labeled with tdTomato. Because all TRAPed, caspase-expressing PAG-USV neurons will subsequently be ablated, viral spread was examined by assessing the absence of tdTomato labeling in the PAG of TRAP2;Ai14 mice.

### Statistics

Parametric, two-sided statistical comparisons were used in all analyses (alpha = 0.05). No statistical methods were used to pre-determine sample size. Same-sex cages of mice were selected at random for inclusion into experimental (PAG-USV_casp_, PAG-USV_GFP_, or PAG-control_casp_). Mice were only excluded from analysis in cases in which viral injections were not targeted accurately. Details of the statistical analyses used in this study are included in Table S1.

### Data availability

All source data generated in this study, as well as custom Matlab codes, will be deposited in a digital data repository, and this section will be modified prior to publication to include the persistent DOI for this dataset.

## Supplemental Information

**Figure S1. Additional quantification of effects of PAG-USV neuronal ablation on female USV production**. Total USVs are plotted for interactions between pairs of females that included (left, N = 12 trials) 1 PAG-USV_GFP_ female and 1 intact female, (middle, N = 12 trials) 1 PAG-USV_casp_ female and 1 intact female, and (right, N = 8 trials) 2 PAG-USV_casp_ females. Data in left and middle columns are the same data represented in Figure 1D, post-4-OHT.

**Figure S2. Female mice produce squeaks during courtship interactions following ablation of PAG-USV neurons**. Blue shading in representative spectrogram indicates squeaks produced by a PAG-USV_casp_ female during an interaction with a male. Courtship USVs produced by the male can also be seen in this spectrogram.

**Table S1. Statistics table**. Details of statistical tests used in the study are provided.

## References

Adametz, J., and O’Leary, J. L. (1959). Experimental mutism resulting from periaqueductal lesions in cats. Neurology 9, 636–636. doi: 10.1212/WNL.9.10.636.

Allen, W. E., DeNardo, L. A., Chen, M. Z., Liu, C. D., Loh, K. M., Fenno, L. E., et al. (2017). Thirst-associated preoptic neurons encode an aversive motivational drive. Science 357, 1149–1155. doi: 10.1126/science.aan6747.

DeNardo, L. A., Liu, C. D., Allen, W. E., Adams, E. L., Friedmann, D., Fu, L., et al. (2019). Temporal Evolution of Cortical Ensembles Promoting Remote Memory Retrieval. Nat Neurosci 22, 460–469. doi: 10.1038/s41593-018-0318-7.

Düsterhöft, F., Häusler, U., and Jürgens, U. (2000). On the search for the vocal pattern generator. A single-unit recording study. Neuroreport 11, 2031–2034. doi: 10.1097/00001756-200006260-00045.

Esposito, A., Demeurisse, G., Alberti, B., and Fabbro, F. (1999). Complete mutism after midbrain periaqueductal gray lesion. Neuroreport 10, 681–685. doi: 10.1097/00001756-199903170-00004.

Gourbal, B. E. F., Barthelemy, M., Petit, G., and Gabrion, C. (2004). Spectrographic analysis of the ultrasonic vocalisations of adult male and female BALB/c mice. Naturwissenschaften 91, 381–385. doi: 10.1007/s00114-004-0543-7.

Grimsley, J. M. S., Hazlett, E. G., and Wenstrup, J. J. (2013). Coding the Meaning of Sounds: Contextual Modulation of Auditory Responses in the Basolateral Amygdala. J Neurosci 33, 17538–17548. doi: 10.1523/JNEUROSCI.2205-13.2013.

Grimsley, J. M. S., Sheth, S., Vallabh, N., Grimsley, C. A., Bhattal, J., Latsko, M., et al. (2016). Contextual Modulation of Vocal Behavior in Mouse: Newly Identified 12 kHz “Mid-Frequency” Vocalization Emitted during Restraint. Front Behav Neurosci 10, 38. doi: 10.3389/fnbeh.2016.00038.

Håkansson, J., Jiang, W., Xue, Q., Zheng, X., Ding, M., Agarwal, A. A., et al. (2022). Aerodynamics and motor control of ultrasonic vocalizations for social communication in mice and rats. BMC Biol 20, 3. doi: 10.1186/s12915-021-01185-z.

Holstege, G. (1989). Anatomical study of the final common pathway for vocalization in the cat. J Comp Neurol 284, 242–252. doi: 10.1002/cne.902840208.

Jürgens, U. (1994). The role of the periaqueductal grey in vocal behaviour. Behav Brain Res 62, 107–117. doi: 10.1016/0166-4328(94)90017-5.

Jürgens, U. (2002). Neural pathways underlying vocal control. Neurosci Biobehav Rev 26, 235–258. doi: 10.1016/S0149-7634(01)00068-9.

Jürgens, U. (2009). The Neural Control of Vocalization in Mammals: A Review. J Voice 23, 1–10. doi: 10.1016/j.jvoice.2007.07.005.

Jürgens, U., and Pratt, R. (1979). Role of the periaqueductal grey in vocal expression of emotion. Brain Res 167, 367–378. doi: 10.1016/0006-8993(79)90830-8.

Kelly, A. H., Beaton, L. E., and Magoun, H. W. (1946). A midbrain mechanism for facio-vocal activity. J Neurophysiol 9, 181–189. doi: 10.1152/jn.1946.9.3.181.

Kittelberger, J. M., Land, B. R., and Bass, A. H. (2006). Midbrain periaqueductal gray and vocal patterning in a teleost fish. J Neurophysiol 96, 71–85. doi: 10.1152/jn.00067.2006.

Larson, C. R., and Kistler, M. K. (1984). Periaqueductal gray neuronal activity associated with laryngeal EMG and vocalization in the awake monkey. Neurosci Lett 46, 261–266. doi: 10.1016/0304-3940(84)90109-5.

Larson, C. R., and Kistler, M. K. (1986). The relationship of periaqueductal gray neurons to vocalization and laryngeal EMG in the behaving monkey. Exp Brain Res 63, 596–606. doi: 10.1007/BF00237482.

Mahrt, E., Agarwal, A., Perkel, D., Portfors, C., and Elemans, C. P. H. (2016). Mice produce ultrasonic vocalizations by intra-laryngeal planar impinging jets. Curr Biol 26, R880–R881. doi: 10.1016/j.cub.2016.08.032.

Moles, A., Costantini, F., Garbugino, L., Zanettini, C., and D’Amato, F. R. (2007). Ultrasonic vocalizations emitted during dyadic interactions in female mice: a possible index of sociability? Behav Brain Res 182, 223–230. doi: 10.1016/j.bbr.2007.01.020.

Neunuebel, J. P., Taylor, A. L., Arthur, B. J., and Egnor, S. E. R. (2015). Female mice ultrasonically interact with males during courtship displays. Elife 4. doi: 10.7554/eLife.06203.

Newman, J. D., and MacLean, P. D. (1982). Effects of tegmental lesions on the isolation call of squirrel monkeys. Brain Res 232, 317–330. doi: 10.1016/0006-8993(82)90276-1.

Nyby, J. (1983). Ultrasonic vocalizations during sex behavior of male house mice (Mus musculus): a description. Behav Neural Biol 39, 128–134. doi: 10.1016/s0163-1047(83)90722-7.

Nyby, J., Wysocki, C. J., Whitney, G., Dizinno, G., and Schneider, J. (1979). Elicitation of male mouse (Mus musculus) ultrasonic vocalizations: I. Urinary cues. J Comp Physiol Psychol 93, 957–975. doi: 10.1037/h0077623.

Riede, T. (2011). Subglottal pressure, tracheal airflow, and intrinsic laryngeal muscle activity during rat ultrasound vocalization. J Neurophysiol 106, 2580–2592. doi: 10.1152/jn.00478.2011.

Riede, T. (2013). Stereotypic laryngeal and respiratory motor patterns generate different call types in rat ultrasound vocalization. J Exp Zool A Ecol Genet Physiol 319, 213–224. doi: 10.1002/jez.1785.

Riede, T., Borgard, H. L., and Pasch, B. (2017). Laryngeal airway reconstruction indicates that rodent ultrasonic vocalizations are produced by an edge-tone mechanism. R Soc Open Sci 4, 170976. doi: 10.1098/rsos.170976.

Roberts, L. H. (1975). The rodent ultrasound production mechanism. Ultrasonics 13, 83–88. doi: 10.1016/0041-624X(75)90052-9.

Ruat, J., Genewsky, A. J., Heinz, D. E., Kaltwasser, S. F., Canteras, N. S., Czisch, M., et al. (2022). Why do mice squeak? Toward a better understanding of defensive vocalization. iScience 25, 104657. doi: 10.1016/j.isci.2022.104657.

Sangiamo, D. T., Warren, M. R., and Neunuebel, J. P. (2020). Ultrasonic signals associated with different types of social behavior of mice. Nat Neurosci 23, 411–422. doi: 10.1038/s41593-020-0584-z.

Shiba, K., Umezaki, T., Zheng, Y., and Miller, A. D. (1997). The nucleus retroambigualis controls laryngeal muscle activity during vocalization in the cat. Exp Brain Res 115, 513–519. doi: 10.1007/pl00005721.

Skultety, F. M. (1962). Experimental mutism in dogs. Arch Neurol 6, 235–241. doi: 10.1001/archneur.1962.00450210063007.

Skultety, F. M. (1965). Mutism in cats with rostral midbrain lesions. 1. Arch Neurol 12, 211–225. doi: 10.1001/archneur.1965.00460260101012.

Tschida, K., Michael, V., Takatoh, J., Han, B.-X., Zhao, S., Sakurai, K., et al. (2019). A Specialized Neural Circuit Gates Social Vocalizations in the Mouse. Neuron 103, 459-472.e4. doi: 10.1016/j.neuron.2019.05.025.

VanderHorst, V. G., Terasawa, E., and Ralston, H. J. (2001). Monosynaptic projections from the nucleus retroambiguus region to laryngeal motoneurons in the rhesus monkey. Neuroscience 107, 117–125. doi: 10.1016/s0306-4522(01)00343-8.

Vanderhorst, V. G., Terasawa, E., Ralston, H. J., and Holstege, G. (2000). Monosynaptic projections from the nucleus retroambiguus to motoneurons supplying the abdominal wall, axial, hindlimb, and pelvic floor muscles in the female rhesus monkey. J Comp Neurol 424, 233–250. doi: 10.1002/1096-9861(20000821)424:2<233::aid-cne4>3.0.co;2-c.

Warren, M. R., Clein, R. S., Spurrier, M. S., Roth, E. D., and Neunuebel, J. P. (2020). Ultrashort-range, high-frequency communication by female mice shapes social interactions. Sci Rep 10, 2637. doi: 10.1038/s41598-020-59418-0.

Warren, M. R., Spurrier, M. S., Roth, E. D., and Neunuebel, J. P. (2018). Sex differences in vocal communication of freely interacting adult mice depend upon behavioral context. PLoS One 13, e0204527. doi: 10.1371/journal.pone.0204527.

Warren, M. R., Spurrier, M. S., Sangiamo, D. T., Clein, R. S., and Neunuebel, J. P. (2021). Mouse vocal emission and acoustic complexity do not scale linearly with the size of a social group. J Exp Biol 224, jeb239814. doi: 10.1242/jeb.239814.

Whitney, G., Alpern, M., Dizinno, G., and Horowitz, G. (1974). Female odors evoke ultrasounds from male mice. Anim Learn Behav 2, 13–18. doi: 10.3758/bf03199109.

Wild, J. M., Kubke, M. F., and Mooney, R. (2009). Avian nucleus retroambigualis: cell types and projections to other respiratory-vocal nuclei in the brain of the zebra finch (Taeniopygia guttata). J Comp Neurol 512, 768–783. doi: 10.1002/cne.21932.

Zhang, S. P., Davis, P. J., Carrive, P., and Bandler, R. (1992). Vocalization and marked pressor effect evoked from the region of the nucleus retroambigualis in the caudal ventrolateral medulla of the cat. Neurosci Lett 140, 103–107. doi: 10.1016/0304-3940(92)90692-z.

Zhao, X., Ziobro, P., Pranic, N. M., Chu, S., Rabinovich, S., Chan, W., et al. (2021). Sex- and context-dependent effects of acute isolation on vocal and non-vocal social behaviors in mice. PLoS One 16, e0255640. doi: 10.1371/journal.pone.0255640.

