## Supplementary material for "Different populations of midbrain neurons control the production of distinct acoustic categories of vocalization": Figs. S1-2, Table S1

### Figure S1. Additional quantification of effects of PAG-USV neuronal ablation on female USV production.

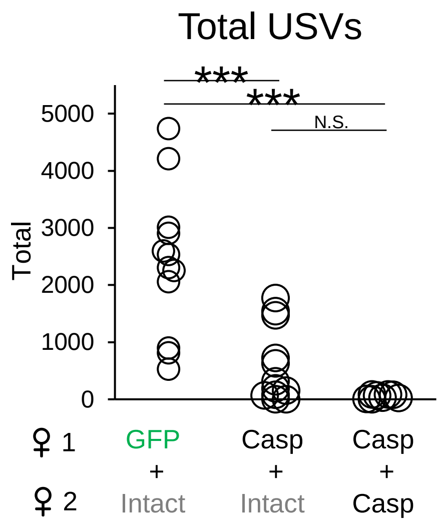

**Figure S1. Additional quantification of effects of PAG-USV neuronal ablation on female USV production.** Total USVs are plotted for interactions between pairs of females that included 1 PAG-$USV_{GFP}$ female and 1 intact female (left, N = 12 trials), 1 PAG-$USV_{casp}$ female and 1 intact female (middle, N = 12 trials) , and 2 PAG-$USV_{casp}$ females (right, N = 8 trials). Please note that data in left and middle columns are the same data represented in Figure 1D, post-4-OHT.

Figure S2. Female mice produce squeaks during courtship interactions following ablation of PAG-USV neurons.

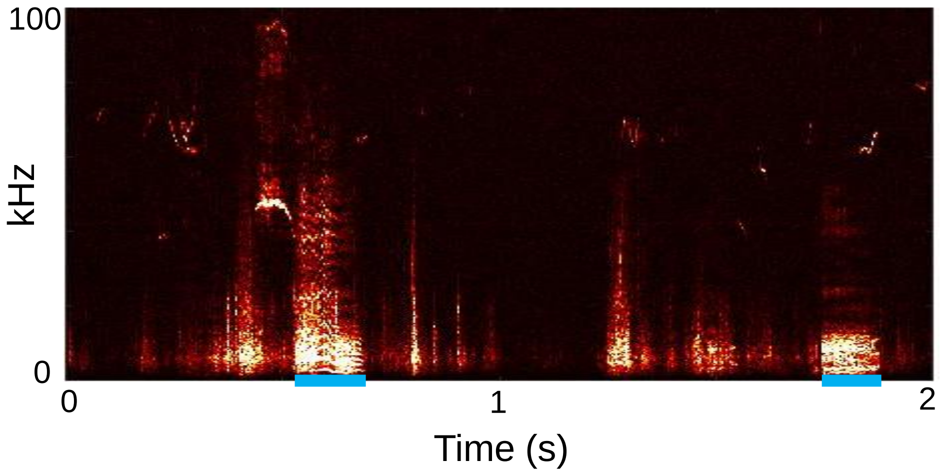

**Figure S2. Female mice produce squeaks during courtship interactions following ablation of PAG-USV neurons.** Blue shading in representative spectrogram indicates squeaks produced by a PAG-USV$_{casp}$ female during an interaction with a male. Courtship USVs produced by the male can also be seen in this spectrogram.

**Table S1**

| **Figure, comparison, and statistical test** | **Group means +/- SD** | **Test results** |
|---|---|---|
| Fig. 1B: total USVs, males<br>• Two-way ANOVA (factor 1 = group, factor 2 = time) with repeated measures on one factor (time); post-hoc Tukey's HSD tests | Pre-4-OHT, $PAG\text{-}USV_{casp}$ males = 1582 ± 928 (N = 11)<br><br>Post-4-OHT, $PAG\text{-}USV_{casp}$ males = 53 ± 84 (N = 11)<br><br>Pre-4-OHT, $PAG\text{-}USV_{GFP}$ males 2309 ± 985 (N = 12)<br><br>Post-4-OHT, $PAG\text{-}USV_{GFP}$ males = 1664 ± 787 (N = 12) | Main effect of group: F(1) = 17.27, **P < 0.001**<br>Main effect of time: F(1) = 38.95, **P < 0.001**<br>Interaction: F(1,21) = 6.66, **P = 0.02**<br><br>Post-hoc pairwise comparisons<br>• $PAG\text{-}USV_{casp}$ pre vs. $PAG\text{-}USV_{casp}$ post: t(21) = 6.179, **P < 0.001**<br>• $PAG\text{-}USV_{casp}$ pre vs. $PAG\text{-}USV_{GFP}$ pre: t(21) = -1.819, P = 0.30<br>• $PAG\text{-}USV_{casp}$ pre vs. $PAG\text{-}USV_{GFP}$ post: t(21) = -0.247, P = 0.99<br>• $PAG\text{-}USV_{casp}$ post vs. $PAG\text{-}USV_{GFP}$ pre: t(21) = 6.921, **P < 0.001**<br>• $PAG\text{-}USV_{casp}$ post vs. $PAG\text{-}USV_{GFP}$ post: t(21) = -9.745, **P < 0.001**<br>• $PAG\text{-}USV_{GFP}$ pre vs. $PAG\text{-}USV_{GFP}$ post: t(21) = -2.724, P = 0.06 |
| Text related to Fig. 1B: change in USV rates<br>• T-test | $PAG\text{-}USV_{casp}$ males = 0.07 ± 0.12 (N = 11)<br><br>$PAG\text{-}USV_{GFP}$ males = 0.71 ± 0.32 (N =12) | t(21) = 6.20, **P < 0.001** |
| Fig. 1C: proportion time spent in resident-initiated interaction, males<br>• Two-way ANOVA (factor 1 = group; factor 2 = time) with repeated measures on one factor (time); post-hoc | Pre-4-OHT, $PAG\text{-}USV_{casp}$ males = 0.25 ± 0.09 (N = 11)<br><br>Post-4-OHT, $PAG\text{-}USV_{casp}$ males = 0.26 ± 0.08 (N = 11)<br><br>Pre-4-OHT, $PAG\text{-}USV_{GFP}$ males = 0.31 ± 0.09 (N = 12) | Main effect of group: F(1) = 0.36, P = 0.56<br>Main effect of time: F(1) = 3.37, P = 0.10<br>Interaction: F(1,21) = 4.87, **P = 0.04**<br><br>Post-hoc pairwise comparisons<br>• $PAG\text{-}USV_{casp}$ pre vs. $PAG\text{-}USV_{casp}$ post: t(21) = -0.324, P = 0.99<br>• $PAG\text{-}USV_{casp}$ pre vs. $PAG\text{-}USV_{GFP}$ pre: t(21) = -1.635, P = 0.38<br>• $PAG\text{-}USV_{casp}$ pre vs. $PAG\text{-}USV_{GFP}$ post: t(21) = -0.538, P = 0.95<br>• $PAG\text{-}USV_{casp}$ post vs. $PAG\text{-}USV_{GFP}$ pre: t(21) = -1.514, P = 0.45 |

| | | |
|---|---|---|
| Tukey's HSD tests | Post-4-OHT, PAG-USV$_{GFP}$ males = 0.24 ± 0.06 (N =12) | • PAG-USV$_{casp}$ post vs. PAG-USV$_{GFP}$ post: t(21) = 0.906, P = 0.80<br>• PAG-USV$_{GFP}$ pre vs. PAG-USV$_{GFP}$ post: t(21) = 2.853, P = 0.04 |
| Fig. 1D: total USVs, females<br>• Two-way ANOVA (factor 1 = group; factor 2 = time) with repeated measures on one factor (time); post-hoc Tukey's HSD tests | Pre-4-OHT, PAG-USV$_{casp}$ females = 1468 ± 599 (N = 12)<br><br>Post-4-OHT, PAG-USV$_{casp}$ females = 579 ± 657 (N = 12)<br><br>Pre-4-OHT, PAG-USV$_{GFP}$ females= 2060 ± 873 (N = 12)<br><br>Post-4-OHT, PAG-USV$_{GFP}$ females = 2410 ± 1273 (N = 12) | Main effect of group: F(1) = 19.11, **P < 0.001**<br>Main effect of time: F(1) = 1.31, P = 0.26<br>Interaction: F(1,22) = 6.93, **P = 0.02**<br><br>Post-hoc pair-wise comparisons<br>• PAG-USV$_{casp}$ pre vs. PAG-USV$_{casp}$ post: t(22) = 2.671, P = 0.06<br>• PAG-USV$_{casp}$ pre vs. PAG-USV$_{GFP}$ pre: t(22) = -1.938, P = 0.24<br>• PAG-USV$_{casp}$ pre vs. PAG-USV$_{GFP}$ post: t(22) = -2.591, P = 0.07<br>• PAG-USV$_{casp}$ post vs. PAG-USV$_{GFP}$ pre: t(22) = 4.073, **P = 0.003**<br>• PAG-USV$_{casp}$ post vs. PAG-USV$_{GFP}$ post: t(22) = -4.428, **P = 0.001**<br>• PAG-USV$_{GFP}$ pre vs. PAG-USV$_{GFP}$ post: t(22) = -1.052, P = 0.72 |
| Fig. 1E: proportion time spent in resident-initiated interaction, females<br>• Two-way ANOVA (factor 1 = group; factor 2 = time) with repeated measures on one factor (time) | Pre-4-OHT, PAG-USV$_{casp}$ females = 0.25 ± 0.05 (N = 12)<br><br>Post-4-OHT, PAG-USV$_{casp}$ females = 0.26 ± 0.05 (N = 12)<br><br>Pre-4-OHT, PAG-USV$_{GFP}$ females = 0.24 ± 0.06 (N = 12)<br><br>Post-4-OHT, PAG-USV$_{GFP}$ females = 0.30 ± 0.10 (N = 12) | Main effect of group: F(1) = 0.71, P = 0.41<br>Main effect of time: F(1) = 1.6, P = 0.22<br>Interaction: F(1,22) = 1.32, P = 0.26 |

| | | |
|---|---|---|
| Fig. 2A: number of footshocks that elicited squeaks (of 10 total)<br>• Two-way ANOVA (factor 1 = sex, factor 2 = group) | PAG-USV$_{casp}$ males = 10.0 ± 0.0 (N = 11)<br><br>PAG-USV$_{GFP}$ males = 9.7 ± 0.9 (N = 12)<br><br>PAG-USV$_{casp}$ females = 10.0 ± 0.0 (N = 12)<br><br>PAG-USV$_{GFP}$ females = 9.9 ± 0.3 (N = 12) | Main effect of sex: F(1) = 0.94, P = 0.34<br>Main effect of time: F(1) = 2.29, P = 0.14<br>Interaction: F(1,43) = 0.76, P = 0.39 |
| Fig. 2B: mean squeak duration<br>• Two-way ANOVA (factor 1 = sex, factor 2 = group) | PAG-USV$_{casp}$ males = 118.8 ± 17.0 (N = 11)<br><br>PAG-USV$_{GFP}$ males = 122.2 ± 20.1 (N = 12)<br><br>PAG-USV$_{casp}$ females = 160.9 ± 41.7 (N = 12)<br><br>PAG-USV$_{GFP}$ females = 142.1 ± 38.1 (N = 12) | Main effect of sex: F(1) = 11.39, **P = 0.002**<br>Main effect of group: F(1) = 0.88, P = 0.41<br>Interaction: F(1,43) = 1.32, P = 0.23 |
| Fig. 2C: mean squeak pitch<br>• Two-way ANOVA (factor 1 = sex, factor 2 = group) | PAG-USV$_{casp}$ males = 5891 ± 333.8 (N = 11)<br><br>PAG-USV$_{GFP}$ males = 6038 545.1 (N = 12)<br><br>PAG-USV$_{casp}$ females = 6141 ± 343.7 (N = 12)<br><br>PAG-USV$_{GFP}$ females = 5884 ± 412.2 (N = 12) | Main effect of sex: F(1) = 0.09, P = 0.75<br>Main effect of group: F(1) =0.43, P = 0.54<br>Interaction: F(1,43) = 2.29, P = 0.14 |
| Fig. 2D: mean squeak amplitude<br>• Two-way ANOVA (factor | PAG-USV$_{casp}$ males = 1.96 ± 0.37 (N = 11)<br><br>PAG-USV$_{GFP}$ males = 2.56 0.31 (N = 12) | Main effect of sex: F(1) = 6.37, **P = 0.01**<br>Main effect of group: F(1) = 16.38, **P < 0.001**<br>Interaction: F(1,43) = 0.83, P = 0.54 |

| | | |
|---|---|---|
| 1 = sex, factor 2 = group) | PAG-USV$_{casp}$ females = 2.37 ± 0.57 (N = 12)<br><br>PAG-USV$_{GFP}$ females = 2.82 ± 0.47 (N = 12) | |
| Fig. 3B: total USVs<br>• Paired t-test | Pre-4-OHT PAG-control$_{casp}$ males = 1596 ± 664 (N = 5)<br><br>Pre-4-OHT PAG-control$_{casp}$ males = 1976 ± 924 (N = 5) | t(4) = -1.71, P = 0.16 |
| Text related to Fig. 3B: mean USV duration<br>• Paired t-test | Pre-4-OHT PAG-control$_{casp}$ males = 47.0 ± 9.1 (N = 5)<br><br>Pre-4-OHT PAG-control$_{casp}$ males = 49.7 ± 9.0 (N = 5) | t(4) = -1.11, P = 0.33 |
| Text related to Fig. 3B: mean USV pitch<br>• Paired t-test | Pre-4-OHT PAG-control$_{casp}$ males = 69225 ± 4380 (N = 5)<br><br>Pre-4-OHT PAG-control$_{casp}$ males = 69773 ± 3328 (N = 5) | t(4) = -0.33, P = 0.76 |
| Text related to Fig. 3B: mean USV amplitude<br>• Paired t-test | Pre-4-OHT PAG-control$_{casp}$ males = 14.9 ± 3.9 (N = 5)<br><br>Pre-4-OHT PAG-control$_{casp}$ males = 11.5 ± 1.0 (N = 5) | t(4) = 2.09, P = 0.11 |
| Fig. 3C: number of footshocks that elicited squeaks (of 10 total)<br>• One-way ANOVA | PAG-control$_{casp}$ males = 9.6 ± 0.5 (N = 5)<br><br>PAG-USV$_{casp}$ males = 10.0 ± 0.0 (N = 11)<br><br>PAG-USV$_{GFP}$ males = 9.7 ± 0.9 (N = 12) | F(2,25) = 1.07, P = 0.36 |

| | | |
|---|---|---|
| Fig. 3D: mean squeak duration<br>• One-way ANOVA | PAG-control$_{casp}$ males = 97.5 ± 20.4 (N = 5)<br><br>PAG-USV$_{casp}$ males = 118.8 ± 17.0 (N = 11)<br><br>PAG-USV$_{GFP}$ males = 122.2 ± 18.5 (N = 12) | F(2,25) = 3.15, P = 0.06 |
| Fig. 3E: mean squeak pitch<br>• One way ANOVA | PAG-control$_{casp}$ males = 6008 ± 242 (N = 5)<br><br>PAG-USV$_{casp}$ males = 5891 ± 334 (N = 11)<br><br>PAG-USV$_{GFP}$ males = 6471 ± 807 (N = 12) | F(2,25) = 3.15, P = 0.06 |
| Fig. 3F: mean squeak amplitude<br>• One way ANOVA with post-hoc Tukey's HSD tests | PAG-control$_{casp}$ males = 0.87 ± 0.25 (N = 5)<br><br>PAG-USV$_{casp}$ males = 1.96 ± 0.97 (N = 11)<br><br>PAG-USV$_{GFP}$ males = 2.56 ± 0.32 (N = 12) | F(2,25) = 48.1, P < 0.001<br><br>Post-hoc pairwise comparisons:<br>• PAG-control$_{casp}$ vs. PAG-USV$_{casp:}$ t(25) = -6.204, **P < 0.001**<br>• PAG-control$_{casp}$ vs. PAG-USV$_{GFP:}$ t(25) = -9.772, **P < 0.001**<br>• PAG-USV$_{casp}$ vs. PAG-USV$_{GFP:}$ t(25) = -4.445, **P < 0.001** |
| Fig. S1: 4-OHT USV rates in female groups<br>• One way ANOVA with post-hoc Tukey's HSD tests | A: Pairs containing 1 PAG-USV$_{GFP}$ female + 1 intact female: 2410 ± 1273 (N = 12)<br><br>B: Pairs containing 1 PAG-USV$_{casp}$ female + 1 intact female: 579 ± 657 (N = 12)<br><br>C: Pairs containing 2 PAG-USV$_{casp}$ females: 49 ± 36 (N = 8) | F(2,29) = 20.96, **P < 0.001**<br><br>Post-hoc pairwise comparisons:<br>• A vs. B**:** t(29) = -5.083, **P < 0.001**<br>• A vs. C**:** t(29) = 5.861, **P < 0.001**<br>• B vs. C: t(29) = 1.315, P = 0.4 |
